# Branching microtubule nucleation is controlled by importin-mediated inhibition of TPX2 phase separation

**DOI:** 10.1101/2020.09.01.276469

**Authors:** Mohammad S. Safari, Matthew R. King, Clifford P. Brangwynne, Sabine Petry

## Abstract

The microtubule-based mitotic spindle is responsible for equally partitioning the genome during each cell division, and its assembly is executed by several microtubule nucleation pathways. In the spindle center, Targeting Protein for XKlp2 (TPX2) promotes branching microtubule nucleation, where new microtubules are nucleated from pre-existing ones. Until the onset of spindle assembly, TPX2 is sequestered by importins-α/β, yet the molecular nature of this regulation remains unclear, particularly since TPX2 was recently found to undergo a liquid-liquid phase separation to execute its function. Here we demonstrate that TPX2 interacts with importins-α/β with nanomolar affinity as a 1:1:1 mono-dispersed trimer. We identify a new nuclear localization sequence (NLS) on TPX2, which contributes to its high-affinity interaction with importin-α. Interestingly, importin-β alone can also associate with TPX2, and does so via dispersed, weak interactions. Interactions of both importin-α and importin-β with TPX2 each inhibit its propensity for phase separation, and consequently its ability to orchestrate branching microtubule nucleation. In sum, our study explains how TPX2 is regulated in order to facilitate spindle assembly, and provides novel insight into how a protein phase separation can be inhibited via weak biomolecular interactions.

**Significance Statement:** The discovery that proteins can undergo phase separation is revolutionizing biology. Characterization of dozens of phase separating proteins in vitro over the past several years has mainly focused on how macromolecules undergo liquid-liquid phase separation (LLPS). The next, and possibly bigger challenge is to investigate how LLPS is regulated in the cell, namely how it is inhibited to spatiotemporally control a certain cellular function. Here, we addressed this challenge by identifying how the spindle assembly factor TPX2 is inhibited by importins from undergoing LLPS and thereby turning on spindle assembly.

## Introduction

The propagation of life requires rapid and accurate assembly of the microtubule (MT)-based mitotic spindle (1, 2). Ran-GTP is a key regulator of both MT nucleation and spindle organization (3, 4), by repurposing the RanGTP-karyopherin nuclear import pathway. Ran gets converted into its GTP-bound state near chromatin and releases spindle assembly factors (SAFs) from sequestration by karyopherins (3, 5). There is a growing repertoire of about two dozen SAFs, but the molecular mechanism of how karyopherins inhibit SAFs, and thereby spindle assembly, is poorly understood (5, 6).

The majority of SAFs are inhibited by the canonical and abundant karyopherin complex, the importin-α/β heterodimer (7, 6, 8, 9, 10). Importin-α contains an importin-β Binding Domain (IBB), which in the absence of importin-β, masks its nuclear localization signal (NLS) binding pocket and prevents its association with NLS-containing proteins. Within the importin-α/β heterodimer, importin-β binds to the IBB of importin-α, thereby making the heterodimer competent to bind the NLS-containing protein (11, 12, 13). Importin-α/β binding to NLS sites on two SAFs has been proposed to sterically block MT binding domains that lie adjacent to the SAF’s NLS (14, 15). However, the possibility of other modes by which importins could inhibit SAFs, has yet to be explored (16, 17).

The SAF and MT-binding protein Targeting Protein for XKlp2 (TPX2) (3) promotes the formation of spindle MTs via branching MT nucleation (18). In this process MTs are autocatalytically amplified from pre-existing ones while preserving their polarity (18, 19). Branching MT nucleation is thought to contribute the majority of spindle MTs (20, 21), and is orchestrated by TPX2 (22, 23). Therefore, TPX2’s inhibition by importin-α/β is critical for the cell cycle and the onset of spindle formation. A fragment of TPX2 that localizes to MTs *in vitro* overlaps with a known importin-α/β binding site (24), which led to the proposal that importin-α/β sterically inhibits MT binding and thereby spindle assembly (14). However, such inhibition of MT localization is neither observed *in vitro* nor in isolated *Xenopus* egg cytosol (12, 25) and the minimal functional fragment of TPX2 for branching MT nucleation does not contain this MT binding region yet can bind to MTs (26). Therefore, we sought to investigate alternative mechanisms by which importin-α/β may inhibit TPX2 function.

Protein phase separation has been implicated in several aspects of spindle assembly (27, 28) (29, 30). Proteins undergo phase separation to achieve a range of cellular functions including compartmentalization, signaling, and reaction enhancement (31). Regulation of macromolecular phase separation and its associated functions is therefore crucial for cellular health. A karyopherin related to importin-α/β (karyopherin 2-β)s was recently shown to prevent aberrant cytoplasmic phase separation of nuclear proteins by engaging in weak associations distributed throughout the karyopherin and target protein (32, 16, 17). We recently uncovered that TPX2 undergoes liquid-liquid phase separation to enhance the reaction efficiency of branching MT nucleation (30). Most importantly, importin-α/β inhibits TPX2 phase separation *in vitro* and TPX2-mediated MT nucleation in isolated *Xenopus* cytosol (30). However, as with many recent studies focusing on biomolecular phase separation, few molecular details are known, and it remains unclear how importins inhibit TPX2 phase separation.

Here, we characterize how importin-α/β interact with TPX2 and determine which interactions are relevant for inhibiting TPX2 phase separation and function. We identified a new NLS on TPX2 for its interaction with importin-α and demonstrate that importin-β inhibits TPX2 condensation by dispersed, weak interactions. As a result, TPX2-mediated branching MT nucleation is inhibited for spindle assembly. These findings highlight a critical role for dispersed, weak interactions in the inhibition of SAFs and spindle assembly by karyopherins, and may also inform how other SAFs are regulated.

## Results

### Importin α or β alone inhibit TPX2-mediated branching MT nucleation

TPX2 induces branching MT nucleation during spindle assembly, which can be directly observed by adding TPX2 to *Xenopus* egg cytosol to induce the formation of branched MT networks (18) (Fig. 1A). Previously, we revealed that the importin-α/β heterodimer inhibits TPX2-mediated branching MT nucleation (30). To decipher the role of individual importins within this inhibition process, we tested whether importin α or β alone can also suppress TPX2-mediated branching MT nucleation. In the importin-α/β heterodimer, importin-β binds to importin-α’s autoinhibitory IBB domain. To mimic this state, we used a truncated form of importin-α, which does not contain the IBB domain (importin-αΔIBB) (11, 24).

**Figure 1.**
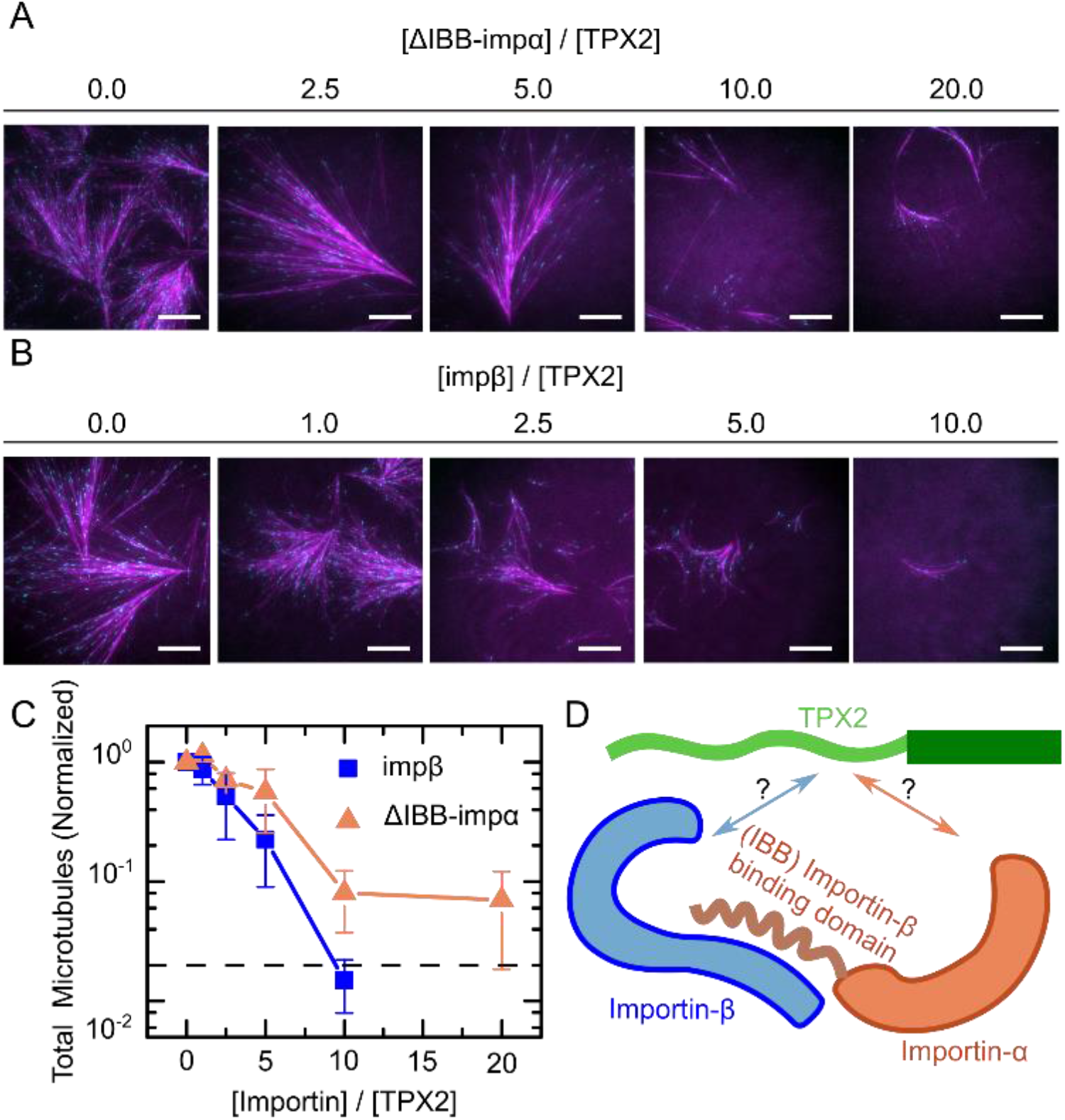
Competency of importins at inhibiting TPX2-mediated branching MT nucleation in cytosol. (*A, B*) Representative TIRF micrographs of TPX2-mediated branching MT nucleation in Xenopus meiotic cytosol in the presence of importin-αΔIBB (*A*) or importin-β (*B*) at indicated fold excess concentrations. Images were taken ~20 minutes into the reaction. Cy5-labeled tubulin (magenta) and mCherry-EB1 (cyan) highlight microtubules and growing microtubule plus ends, respectively. (*C*) Total number of MTs generated by 20 minutes at indicated fold-excess concentration of importin-αΔIBB (dark orange) or importin-β (blue). Each data point represents total number of MTs in an analyzed field of view, which is normalized to the no importin condition from that experimental set. The dashed line represents the extract background activity. (*D*) Schematic of TPX2:importin-α/β heterodimer. In the importin-α/β complex, the Nuclear Localization Sequence (NLS) binding pockets of importin-α are exposed upon binding of importin-β to the IBB (importin-β Binding) domain of importin-α (a.a. 1-70).

When TPX2 was added to *Xenopus* egg cytosol with only importin-αΔIBB at ≥10-fold molar excess, the total number of MTs was drastically reduced (Fig. 1A, C). Similarly, TPX2 addition with only importin-β at 10-fold excess an almost complete reduction in MT number, thus completely inhibiting branching MT nucleation (Fig. 1B-C). These data show that, surprisingly, importin-α and importin-β can each independently inhibit TPX2-mediated branching MT nucleation, with importin-β being particularly effective.

### Importin α or β alone inhibit TPX2 condensation

Recently, we showed that TPX2 forms a co-condensate with tubulin on the MT lattice, which enhances reaction kinetics of branching MT nucleation (30). Moreover, TPX2 binding to MTs is the first essential step to build a branch site by recruiting additional branching factors (22, 23). Thus, the ability to inhibit TPX2 phase separation could be key to the regulation of branching MT nucleation, and the onset of spindle assembly. Therefore, we investigated how importin-α/β affects TPX2 phase separation.

We monitored TPX2 condensation via fluorescence microscopy upon addition of either importin. Addition of importin-αΔIBB to TPX2 resulted first in an enhancement of TPX2 condensation at lower concentrations followed by an inhibition of TPX2 condensation at around 16-20-fold excess (Fig. 2A). In contrast, importin-β and importin-α/β were more effective, and inhibited TPX2 condensation at 5-6 and 2-3-fold excess, respectively (Fig. 2B-C).

**Figure 2.**
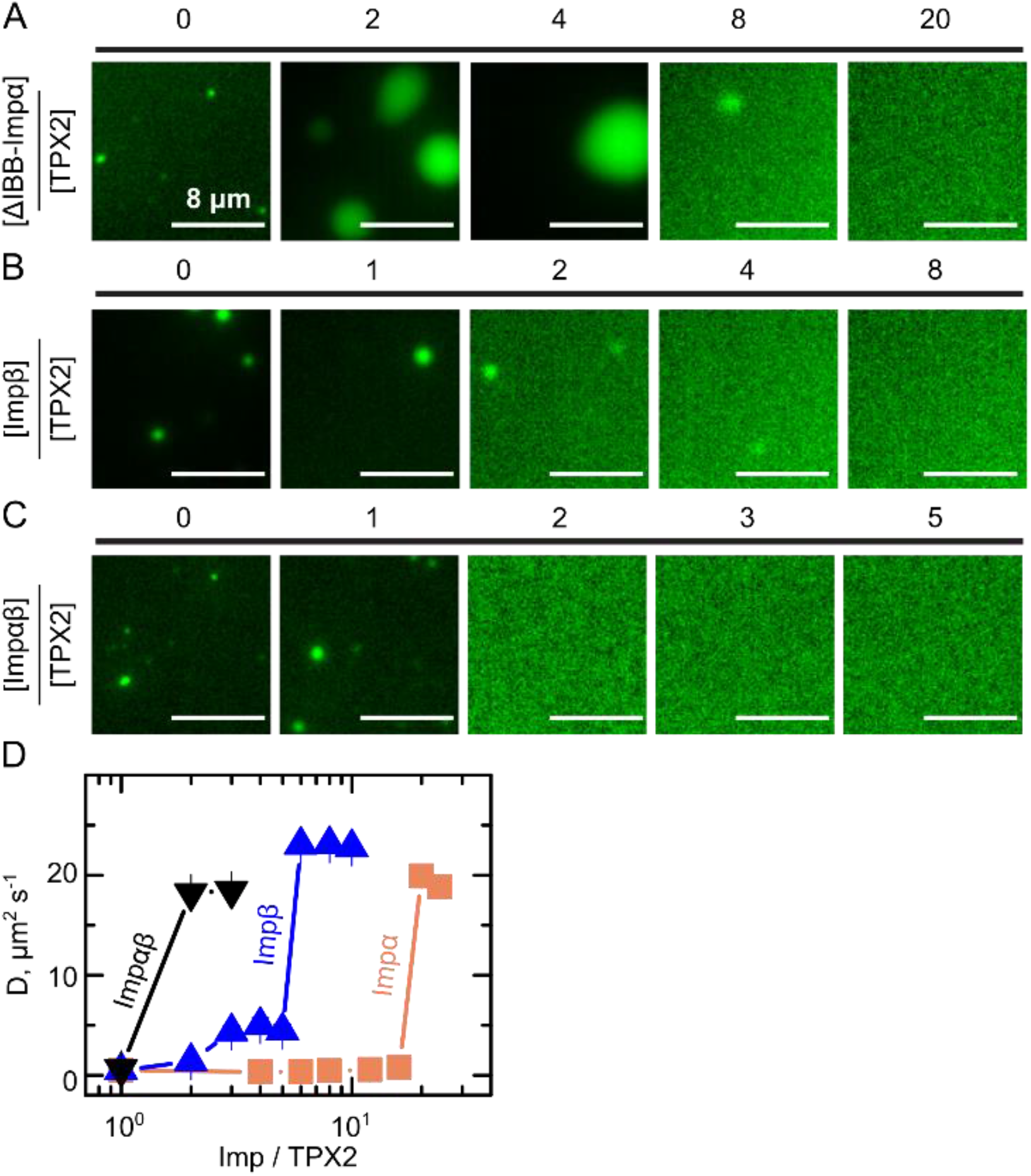
Competency of importins at inhibiting TPX2 condensation. (*A-C*) Representative epifluorescence microscopy images of TPX2 in the presence of (*A*) importin-αΔIBB, (*B*) importin-β, and (*C*) importin-α/β at indicated excess ratio of importin to TPX2. TPX2 concentration is fixed at 1μM. (*D*) Diffusion coefficient of TPX2 solutions in CSF buffer (+ 0.1 M KCl) in the presence of indicated ratio of importin-αΔIBB, importin-β, or importin-α/β to TPX2 concentration as measured by dynamic light scattering. The error bars are calculated from ten distinct intensity-intensity correlation functions. TPX2 concentration is fixed at 1 μM.

To ensure that importin-inhibited TPX2 solutions are indeed monodisperse and they do not contain any sub-resolution condensate, we tested each sample with dynamic light scattering, which measures an intensity-intensity autocorrelation function of scattered light from the solution. Because the scattered intensity is proportional to sixth power of particle sizes, any residual amount of condensate can be detected, in the form of a very small average diffusion coefficient. Our light scattering data further validates the results obtained by light microscopy: the competency of importins in abrogating TPX2 condensation follows importin-α/β > importin-β > importin-αΔIBB (Fig. 2D). Thus, importin-β suppresses TPX2 phase separation even more efficiently than importin-αΔIBB, which harbors the known TPX2 binding sites; this parallels our findings that importin-β inhibits branching MT nucleation more strongly than importin-αΔIBB (Fig. 1A,B,C).

### TPX2 strongly associates with importin-α/β to form a trimer

In order to understand their ability to suppress TPX2 condensation and function in MT branching nucleation, we next investigated how the importin-α/β heterodimer interacts with TPX2. While small fragments of TXP2 has been shown to bind to importin-αΔIBB (24, 12), it remains to be tested how full-length TPX2 associates with the importin-α/β heterodimer (Fig. 1D). We first addressed this question by determining the molecular weight and stoichiometry of the TPX2-importin-α/β complex via size exclusion chromatography in line with multi-angle light scattering (SEC-MALS) (Fig. 3A). GFP-TPX2 eluted off the SEC column with an average molecular weight of 159 ± 52 kDa, comparable to its predicted molecular weight of 112.1 kDa, indicating that it exists mainly as a monomer with no detectable aggregate/oligomers (Fig. 3A-green/dotted curve and Fig. S1A).

**Figure 3.**
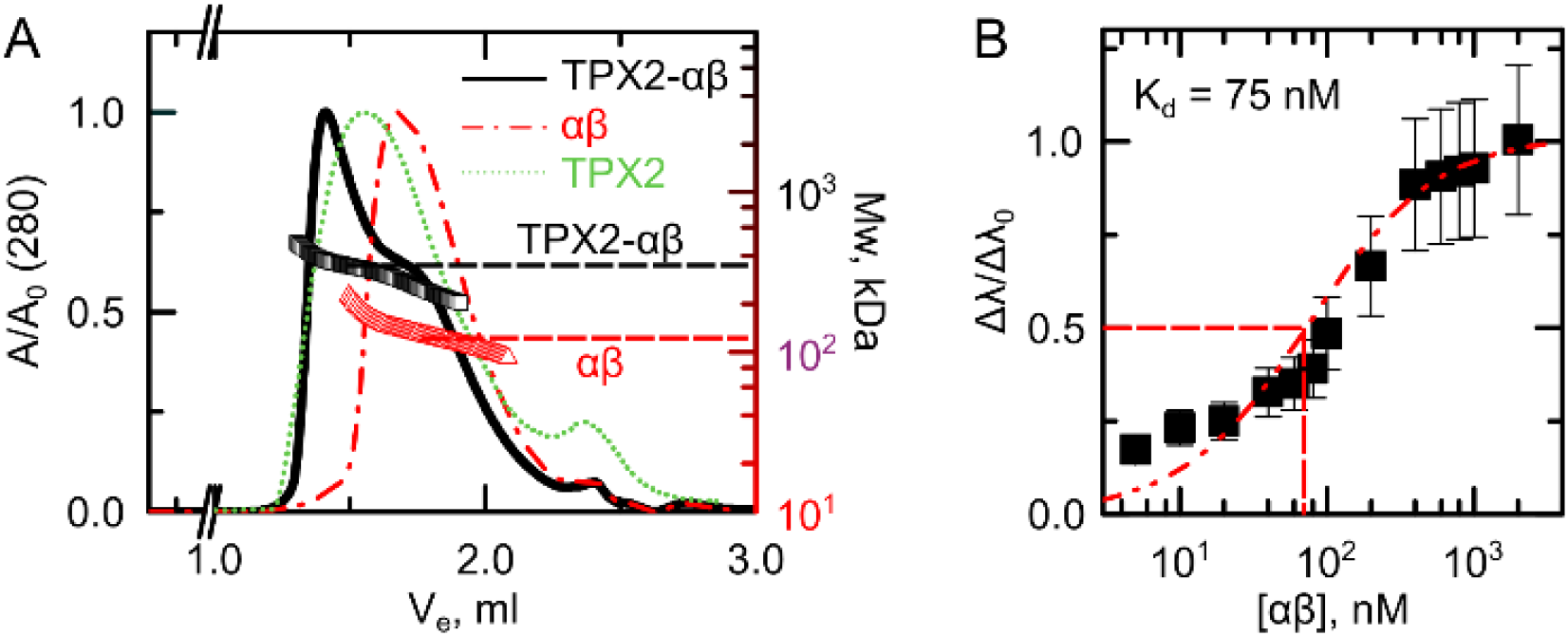
TPX2 strongly associates with importin-α/β to form a trimer. (*A*) Size exclusion chromatography in line with light scattering (SEC-MALS) reveals a stoichiometry of 1:1 for importin-α/β and 1:1:1 binding for TPX2-importin-α/β trimer. The left axis is normalized absorbance (λ = 280 nm) and the right axis shows the molecular weight of eluted complexes. The molecular weight of eluted complex for TPX2:importin-α/β and importin-α/β is shown with black (310 kDa) and red (130 kD) arrows respectively. TPX2 and importin-α/β concentrations are 5 μM and 15 μM, respectively. (*B*) Biolayer interferometry (Octet) normalized amplitude as a function of importin-α/β concentration. The measured binding constant is K_d_ = 75 ± 15 nM. The error bars re calculated from two distinct measurements.

We then allowed GST-importin-α to bind to importin-β and assessed formation of the importin-α/β heterodimer via SEC-MALS. The eluted complex exhibited a molecular weight of 130.0 ± 29 kDa determined via MALS, consistent with the predicted molecular weight of the importin-α/β heterodimer in a 1:1 stoichiometry (165.0 kDa) (Fig. 3A-red/dashed curve and Fig. S1B).

When TPX2 and importins-α/β were allowed to bind, they eluted as a complex faster than TPX2 and importins-α/β alone. This rapid elution reflects a larger molecular weight of 310 ± 70 kDa via MALS (Fig. 3A, black/solid curve) similar to the predicted molecular weight of 286.8 kDa. This suggest that TPX2-importin-α/β exists as a trimer with 1:1:1 stoichiometry.

In order to determine how strongly TPX2 interacts with importin-α/β, we determined their equilibrium dissociation constant. We utilized bio-layer interferometry, which is based on refractive index fluctuations upon binding and dissociation events of target proteins on the sensor surface (33, 13). We determined a K_d_ between TPX2 and importin-α/β of 75 ± 15 nM (Fig. 3B), indicating that TPX2 binds strongly to importin-α/β.

### TPX2 interacts strongly with importin-α via nuclear localization sequences

To address how importin-α and importin-β alone can inhibit TPX2, we examined how each individual importin binds to TPX2 and contributes to forming the TPX2-importin-α/β trimer. By applying biolayer interferometry, we measured a dissociation constant of K_d_ = 61 ± 10 nM between importin-αΔIBB and TPX2 (Fig. 4B), indicating it alone has a strong affinity for TPX2.

**Figure 4.**
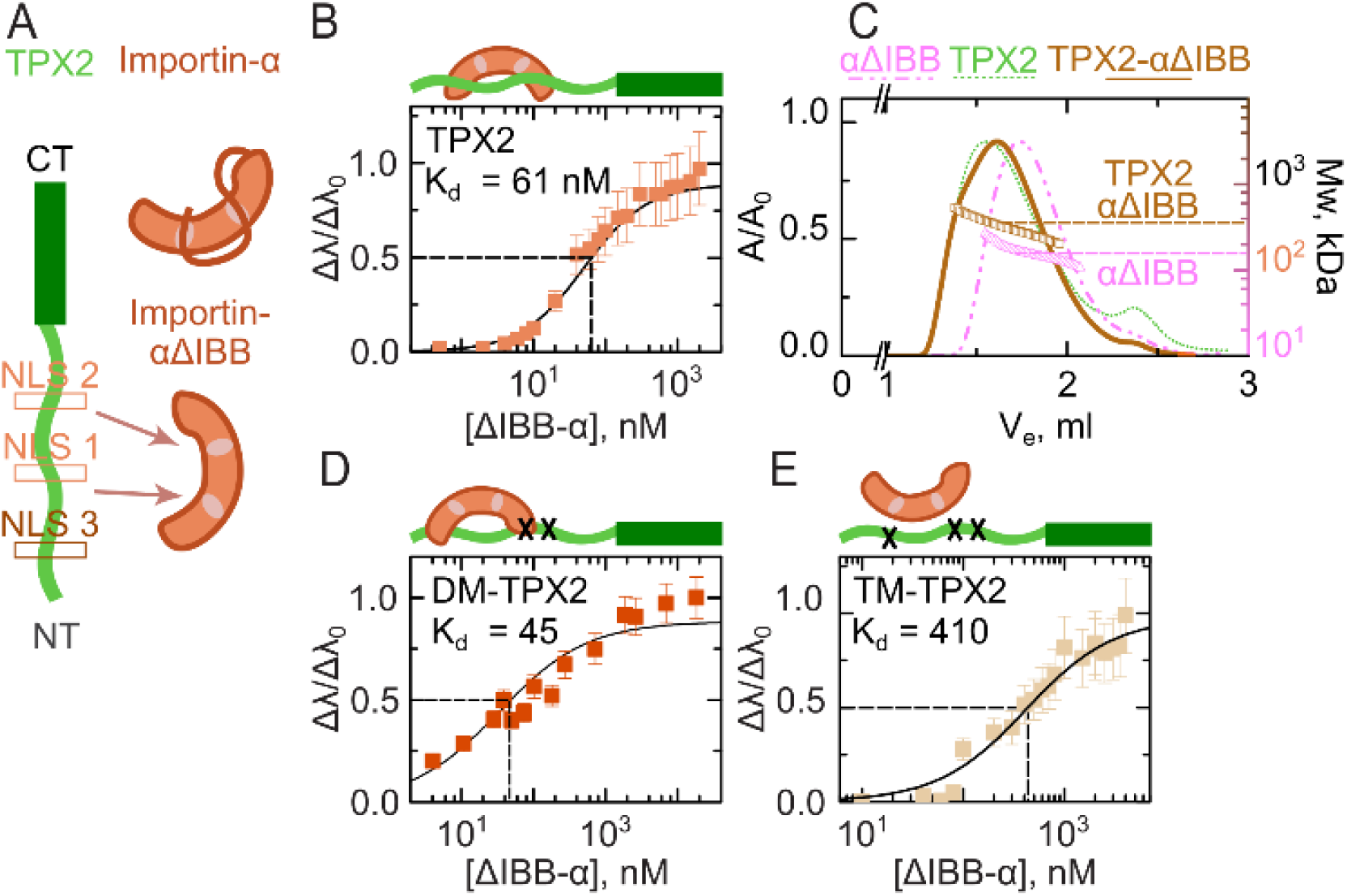
TPX2 interacts with importin-αΔIBB via nuclear localization sequences at a.a. 123 and a.a. 284. (*A*) The architecture of TPX2, importin-α and importin-αΔIBB. TPX2 is comprised of a disordered N-terminal region (light green – a.a.1-480) and an ordered C-terminal region (dark green - a.a. 480-716). The two previously reported nuclear localization sequences (NLSs) are shown in light brown, centered at NL1 a.a. 284 and NL2 a.a. 327. The newly identified putative NLS3 is shown in brown, centered at a.a. 124. Importin-α, in the absence of importin-β, exists in an autoinhibited conformation wherein the NLS binding pockets are occluded by the IBB domain. A truncated version of importin-α without the IBB domain (importin-αΔIBB) has exposed NLS binding pockets. (*B*) TPX2 binding to importin-αΔIBB is mediated by NLS’s at a.a. 124 and a.a. 284. Biolayer interferometry (Octet) of wild-type TPX2 with importin-αΔIBB. Wild-type TPX2 and importin-αΔIBB associate strongly with K_d_ = 61 ± 10 nM. (*C*) TPX2 interacts with importin-αΔIBB as dimer/trimer complex. Size exclusion chromatography in line with light scattering (SEC-MALS) reveals a stoichiometric 2:1/1:2 and 1:1 binding for importin-αΔIBB with TPX2. The left axis is normalized absorbance (λ = 280 nm) and the right axis shows the molecular weight of the eluted complex. The TPX2 and importin-αΔIBB concentrations are 2 μM and 20 μM, respectively. (*D-E*) Biolayer interferometry (Octet) of double mutant TPX2 (DM-TPX2) and triple-mutant TPX2 (TM-TPX2) with importin-αΔIBB, respectively. DM-TPX2 for which the two NLSs (lysines at 284/5/7, and 325/8 are mutated to alanines) and importin-αΔIBB associate strongly with K_d_ = 45 ± 6 nM. Triplemutant TPX2 (TM-TPX2), for which the three NLSs (lysines at 123/4/6, 284/5/7, and 325/8 are mutated to alanines) exhibits a 10-fold lower binding affinity to importin-αΔIBB, K_d_ = 410 ± 70 nM. Values are normalized amplitude as a function of importin-αΔIBB concentration. Error bars are calculated from two distinct measurement.

We next asked whether this interaction is mediated by the two nuclear localization sequences NLS1 and NLS2, which had previously been shown to bind to importin-α fragments (Fig. 4A) (24). We created a double-mutant TPX2 (DM-TPX2), in which NLS1 and NLS2 were mutated to alanines at key residues (in NLS1, K284A, R285A, and in NLS2, K327A, and K330A were mutated). Surprisingly, the DM-TPX2 exhibited a similar binding affinity to importin-αΔIBB, with a dissociation constant K_d_ =45 ± 6 nM (Fig. 4D). This suggests that another strong binding site on TPX2 must exist to allow importin-αΔIBB to bind.

To investigate where this potential new binding site is located, we assessed whether it lies within the C-terminal half of TPX2 (amino acids 319-716). However, this construct exhibited extremely weak association with importin-αΔIBB (Fig. S2), suggesting that an additional binding site must exist within TPX2’s N-terminal half (amino acids 1 – 318).

Since NLS1 and NLS2 were previously identified by scanning TPX2 amino acids 270 to 350 (24), we focused the search for a new binding site to amino acids 1 to 260 of TPX2. We created three TPX2 constructs that cover amino acids 1-99, 1-178, and 1-260. While the short construct 1-99 exhibited only weak binding to importin-αΔIBB, the two longer TPX2 constructs 1-178 and 1-260 exhibited strong binding with dissociation constants of 15 ± 3 and 20 ± 4 nM, respectively (Fig. S2). Bioinformatic analysis revealed a putative NLS sequence within TPX2, in amino acids KKLK located at positions 123-126. To test whether this sequence indeed functions as an NLS, we mutated the lysines to alanines within this motif of TPX2 1-178 (K123A, K124A, K126A), which resulted in a 10-fold loss of binding to importin-αΔIBB compared to the wild type-1-178 TPX2 construct (Fig. S2). This indicates that the KKLK region (amino acids 123-126) of TPX2 constitutes another NLS sequence, which facilitates binding to importin-αΔIBB, and which we therefore term NLS3 (Fig. S2).

To assess whether NLS1, NLS2 and NLS3 are indeed the major and only bindings sites on TPX2 for importin-αΔIBB, we created a triple-mutant TPX2 (TM-TPX2), for which all the NLS sequences were modified (K123A, K124A, K126A, K284A, R285A, K327A, and K330A). Strikingly, TM-TPX2 exhibited a 10-fold decrease (K_d_ = 410 ± 70 nM) in the strength of its interaction with importin-αΔIBB, compared to FL-TPX2 (Fig. 4B, D), suggesting that these three represent all NLS sites that promote specific interactions of TPX2 with importin-αΔIBB.

Because TPX2 contains three NLS sequences to which importin-αΔIBB can bind, we investigated the stoichiometry of the TPX2-importin-αΔIBB complex via SEC-MALS. The eluted complexes exhibited molecular weight of 326 kDa with dispersity of ±88 kDa (Fig. 4C). Comparison of the TPX2-importin-αΔIBB MALS profile to importin-αΔIBB alone, which displays a molecular weight of 159 kDa implying it dimerizes, suggests TPX2:importin-αΔIBB is relatively dispersed and exist in a complex with stoichiometries above 1:1 (Fig. 4C). Taken together, these data show that importin-αΔIBB strongly interacts with TPX2 via three NLS sequences.

### TPX2 weakly and reversibly associates with importin-β

Importin-β is known to associate with nuclear proteins (8) and affect spindle assembly (34) independent of importin-α. Given that importin-β can strongly suppress both TPX2 phase separation and TPX2-mediated branching nucleation (Fig 1,2), we reasoned that importin-β could associate with TPX2 in the absence of importin-α, rather than solely acting as an inert adaptor to occlude the IBB domain of importin-α, according to the existing model. We tested the binding affinity between TPX2 and importin-β by biolayer interferometry. Surprisingly, importin-β and TPX2 only weakly associate, with a dissociation constant of Kd=530 ± 75 nM. Interestingly, this association is driven by TPX2’s N-terminus (a.a. 1-480), which exhibits a similar dissociation constant of 512 ± 70 nM (Fig. 5A), while TPX2’s C-terminus (a.a. 480-716) displays almost no binding with a K_d_ > 4 μM (Fig. S1D).

**Figure 5.**
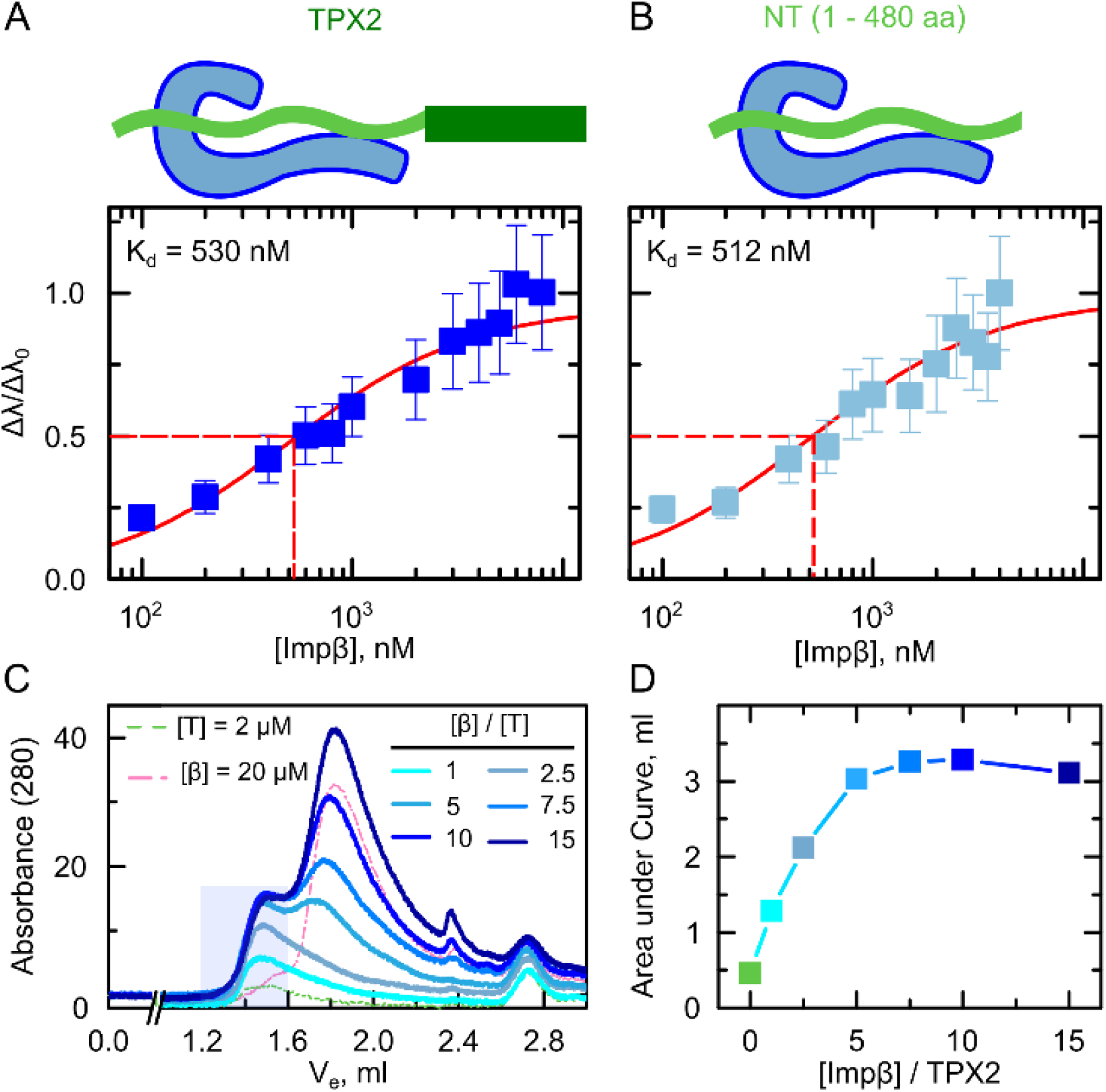
TPX2 weakly and reversibly associates with importin-β. (*A-B*) Importin-β weakly associates with TPX2-N-terminus. Biolayer interferometry (Octet) measuring the association of importin-β with (*A*) full length TPX2, K_d_ = 530 ± 75 nM, and (*B*) the N-terminal region (a.a. 1-480) of TPX2, K_d_ = 512 ± 70 nM. Values are normalized amplitude as a function of importin-β concentration. Error bars are calculated from two distinct measurements. (*C-D*) TPX2-importin-β complex is reversible. (*C*) Size exclusion chromatography with TPX2:importin-β complex exhibits dissolution of the complex upon varying the importin-β concentration under constant TPX2 concentration. The left axis is absorbance (λ = 280 nm). The dashed green and pink chromatograms are TPX2 and importin-β alone at concentrations 2 μM and 20 μM, respectively. (*D*) Area under the curve in the complex elution area (1.2 – 1.6 ml) exhibits an equilibrium saturation for varying concentrations of importin-β.

Given the surprisingly weak interactions between TPX2 and importin-β, we next examined the stoichiometry of the TPX2-importin-β complex. The SEC-MALS data displayed complexes eluting in the range of 200-700 kDa, indicative of a range of oligomeric species (Fig. S1C), which could be due to reversible assembly or irreversible aggregation. To test this hypothesis, we conducted a series of size exclusion chromatography experiments keeping TPX2 at 2 μM and varying importin-β concentration ranging from 2 to 30 μM, and integrated the area under the peak corresponding to the eluted complexes from SEC-MALS (blue shading-Fig. 5C). If these were irreversible aggregates, a linear dependence on Importin-β concentration would be expected. If there was no interaction between importin-β and TPX2, the integrated area should exhibit a constant line with no dependence on Importin-β concentration. Interestingly, we observed a sigmoidal binding curve (Fig. 5D) suggesting that there exists a dynamic equilibrium which saturates upon addition of importin-β at 3-fold excess. To confirm that the saturation is not due to consumption of TPX2, we calculated the un-bound fraction of TPX2 using dissociation constant of K_d_ = 0.5 μM measured by our BLI method. At the maximum concentration of Importin-β, c = 30 μM, more than 5% of TPX2 is still in the un-bound state. In order to completely drive TPX2 to the bound state, the Importin-β concentration must exceed 100 μM. Therefore, the saturation of SEC curve was not due to the depletion of TPX2 and rather due to equilibrium nature of TPX2: Importin-β complex. Thus, our data demonstrate that TPX2 weakly and reversibly associates with importin-β.

### Why is importin-β effective in inhibiting TPX2?

To address this question, we quantified the osmotic compressibility of importin solutions via the second virial coefficient (B_2_) obtained from static light scattering. The low pI of importins (pI < 5.5) and the high negative surface charge suggests that the proteins should be self-repulsive (Fig. S3A), which would result in a positive B_2_ (35). Strikingly, importins exhibited a negative B_2_ (slope in Fig. S3C), indicative of strong attractive intermolecular van der Waals forces originated from the exposed hydrophobic residues on the surface of importins (Fig. S3B) (36, 37). Interestingly, the magnitude of B2, and therefore the strength of intermolecular van der Waals forces, follows the competency of importins to impede the condensates, i.e. |*B*_2_|: importin-α/β > importin-β > importin-αΔIBB. This observation suggests an important role for the short-range van der Waals forces in tuning TPX2 condensation. Moreover, the weak nature of van der Waals interactions could explain why excess molar amounts of importins were required to inhibit TPX2 condensation, similar to the excess molar of Kr2B needed to inhibit FUS condensation (16, 32).

## Discussion

Here we investigated the mechanism of TPX2 regulation by importins-α/β. Previous structural studies indicate that inhibition of TPX2 function may be achieved by importin-α directly blocking a MT binding region of TPX2 (14, 24). While elegant, this mechanism appears to be incomplete since importin-α/β do not block TPX2 localization to MTs *in vitro* or in cytosol (25) and the MT nucleation function of TPX2 resides in a C-terminal region of TPX2 that lacks the putative MT localization domain (18).

The canonical molecular architecture of a 1:1:1 trimer of NLS-protein: importin-α: importin-β has been inferred from many studies, such as for the classic nuclear protein nucleoplasmin (24) and a truncation of the SAF NuMA (15), but not for a full-length SAF. It was not evident *a priori,* that TPX2 would engage with importin-α/β as a trimer as predicted, since TPX2 is ~4x longer, mostly disordered (≥70%) (26), and associates non-stoichiometrically with other interaction partners (38, 30). Nonetheless, our data indicates that the three proteins form a stable, 1:1:1 trimer via high affinity NLS interactions (Fig. 3).

We investigated the nature of intermolecular forces driving the formation of the TPX2:importin-α/β tri-complex by determining the affinity and stoichiometry with which individual importins interact with TPX2. Importin-αΔIBB was previously shown to bind to TPX2 via NLS sequence located at a.a. 284-287 (24, 12). While the a.a. 284-287 NLS is well established, mutating it does not abrogate nuclear import of TPX2 (39). Using TPX2 truncations and mutants, we discovered the existence of an NLS sequence located at a.a. 123-126 of TPX2 (KKLK). This new NLS can mediate TPX2-importin-αΔIBB complex association in the absence of the a.a. 284-287 NLS (Fig. 4). These data suggest that the three NLSs on TPX2 function redundantly to ensure the regulation of TPX2. This is consistent with its demonstrated role in nuclear import, where a co-mutation with a putative NLS at a.a. 123-126 leads to a more significant reduction in nuclear transport (39).

Canonically, nuclear proteins associate with importin-α/β exclusively through an NLS-importin-α interaction (11). Some proteins, however, functionally interact with importin-β either alone or when it is in complex with importin-α (8). Furthermore, some SAFs are thought to exclusively interact with importin-β (6, 9) and importin-β alone can inhibit Ran-GTP mediated MT nucleation (34). Nonetheless, biochemical details of SAF-importin-β interactions remain uninvestigated. Our data demonstrate that importin-β associates with TPX2 via weak interactions, which is reminiscent of other disordered proteins with a different karyopherin-Transportin-1/Karyoperhin-2β (16, 17, 32). Moreover, our data reveals that these weak interactions between TPX2 and importin-β are reversible and promote equilibrium assemblies rather than irreversible aggregation (Fig. 5). In sum, our investigations reveal molecular insight into the architecture of the TPX2:importin-β complex, which is based on dispersed, weak, reversible interactions. While such interactions have been implicated in driving condensation (40, 41, 42, 43, 44), we show here that dispersed, weak, reversible interactions are also implicated in preventing LLPS.

Interestingly, inhibition of TPX2 condensation, and also inhibition of branching MT nucleation, by importins displayed a trend opposite to affinity measurements (K_d_ values). Specifically, the hierarchy of affinity is importin-αΔIBB ≈ importin-α/β >> importin-β, whereas the hierarchy of inhibition competency is importin-α/β > importin-β >> importin-αΔIBB (Figs. 1,2). Thus, weak interactions can be a driving force to inhibit LLPS.

Finally, due to higher competency of importin-β to inhibit TPX2 condensation compared to importin-αΔIBB, and the ability of importin-β to solely interact with TPX2-NT, we speculate that importin-α acts as a bridge to bring importin-β in proximity of TPX2, which can further suppress the TPX2-NT intermolecular forces and abolish the condensation process (Fig 6). In summary, our study highlights how protein phase-separation is inhibited in order to regulate the onset of spindle assembly.

**Figure 6.**
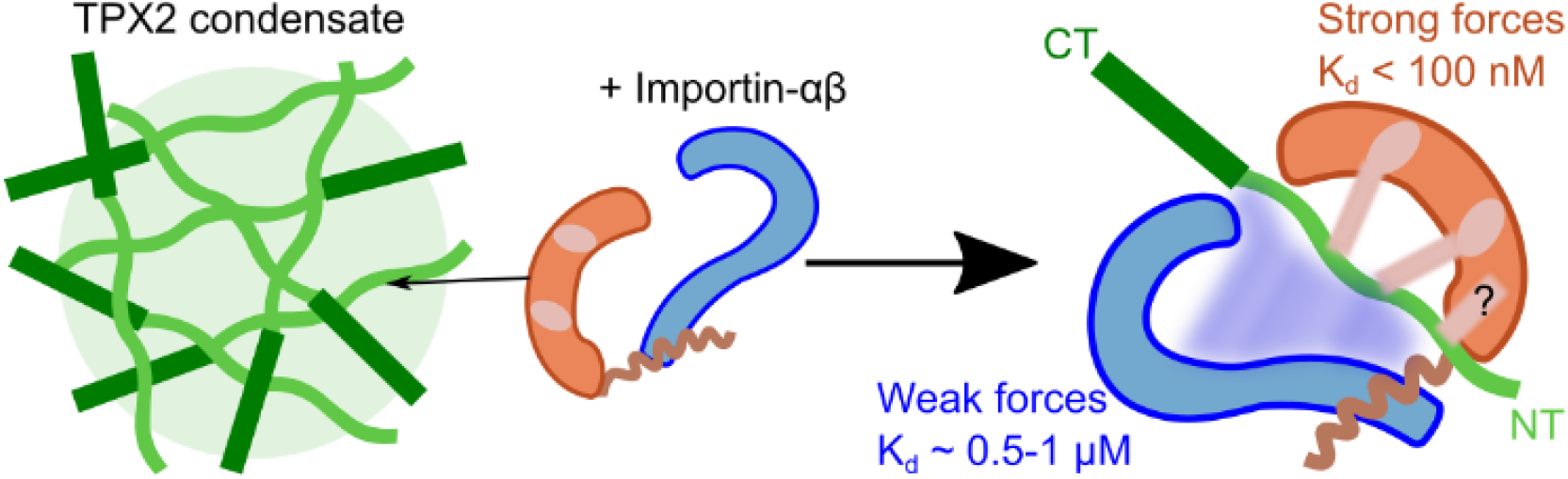
Schematic of TPX2:importin-α/β tri-complex. Importin-α association with TPX2 is mediated via nuclear localization signals of TPX2, which brings importin-β in proximity of TPX2. Importin-β interacts with TPX2-NT via weak and dispersed forces. The precise domain of importin-α to which NLS3 can bind to be determined (?).

## Materials and Methods

Please see supplemental information for details. Recombinant proteins are all *Xenopus laevis* versions and were purified to >95% purity. Size-Exclusion Chromatography (SEC) was carried out on an AKTA Pure-25L. SEC in line with Multi-Angle Light Scattering (SEC-MALS) was done with a 3.2/300 column packed with Superdex-200-increase-in line with a Wyatt light scattering machine. Biolayer interferometry was carried out by varying importin concentration and measuring binding to TPX2 that was attached to sensors, according to vendor specifications of ForteBio. Condensation (phase separation) was achieved by diluting protein mixtures at 0.5M into 0.1M salt (KCl). Static and Dynamic Light Scattering (SLS and DLS) measurements were taken at fixed right angle. Normalized intensity-intensity correlation functions of 20 seconds duration were collected. To visualize TPX2-mediated MT nucleation at indicated excess molar ratios of importins, naturally meiotically arrested *Xenopus laevis* egg cytosol was immunodepleted of endogenous TPX2, indicated proteins were added and the reaction was imaged over time. Images in each panel are representative crops from a single experimental set adjusted to optimal brightness to allow visualization of the relevant structures.

## Acknowledgments

We thank members of the Petry Lab for support of this work and discussions. We thank professor Peter. G. Vekilov for the edits to discussion of the paper, and Dr. Venu Vandavasi for discussion of the interaction studies. This work was supported by Ph.D. training grant T32GM007388 by NIGMS of the National Institutes of Health (to M.R. King), as well as the New Innovator Award of NIGMS of the National Institutes of Health (DP2), the Pew Scholars Program in the Biomedical Sciences, and the David and Lucile Packard Foundation (all to S. Petry). The Brangwynne lab is supported by the Howard Hughes Medical Institute.

## Supplementary Information

### Supplementary Information Text

#### Materials and Methods

| Proteins used |  |  |
| --- | --- | --- |
| Name | Details | Source |
| TPX2 | StrHisGFP-TEV-FL_TPX2 (Full Length - aa1-716) | King MR and Petry S., 2020 |
| ΔNLS-TPX2 | StrHisGFP-TEV-TM_TPX2 (aa1-716 with K123A, K125A, K126A, K284A, K285A, K327A, K330A) | This paper |
| TPX2-DM | StrHisGFP-TEV-DM_TPX2 (aa1-716 with K284A, K285A, K327A, K330A) | This paper |
| TPX2-NT | StrHisGFP-TEV-NT_1-480_TPX2 (N-Terminal aa1-480) | King MR and Petry S., 2020 |
| TPX2-CT | StrHisGFP-TEV-CT_480-716_TPX2 (C-Terminal aa480-716) | King MR and Petry S., 2020 |
| TPX2 aa 1-260 | StrHisGFP-TEV-NT_1-260_TPX2 (N-Terminal aa1-260) | This paper |
| TPX2 aa 1-178 | StrHisGFP-TEV-NT_1-178_TPX2 (N-Terminal aa1-178) | This paper |
| TPX2 aa 1-99 | StrHisGFP-TEV-NT_1-99_TPX2 (N-Terminal aa1-99) | This paper |
| TPX2 aa 1-178Δ123 | StrHisGFP-TEV-NT_1-178_TPX2 (N-Terminal aa1-178, with K123A, K125A, K126A) | This paper |
| TPX2 aa 319-716 | StrHisGFP-TEV-CT_480-716_TPX2 (C-Terminal aa319-716) | King MR and Petry S., 2020 |
| importin-β | importin-β (from cleaved from His-GFP-PreScission-importin-β) | This paper |
| importin-α | GST-PreScission-importin-α | This paper |
| importin-$\alpha\Delta$IBB | GST-PreScission-importin-α (aa71-end -importin-β Binding domain (IBB) removed) | This paper |
| His-mCherry-EB1 | His-mCherry-End Binding Protein 1 | Thalwani et al., 2018 |
| Cy5-Tubulin | NHS-linked Tubulin (Source: PurSolutions LLC: 032005) | Thalwani et al., 2018 |
| BSA | Bovine Serum Albumin | Fisher: 23209 |

##### *E. Coli* strains and growth conditions

DH5α cells (New England Biolabs (NEB):C2987I) were used for all subcloning steps. Rosseta2 cells (Fisher: 71-403-4) were used to express proteins for purification. Cells were grown at various volumes in LB Broth (Sigma: L3522), and cells containing importin-α/β constructs were grown in TB media (Sigma: T0918) prepared according to supplier’s instructions. For protein expression, constructs were transformed into Rosseta2 *E. coli* cells and were grown in temperature-controlled incubators shaking at 180 RPM. Cells were grown at 37°C (0.5-0.7 OD_600_), and induced with 0.75 mM isopropyl-β–D-1-thiogalactopyranoside (IPTG) for another 6 hours at 25°C. Cell pellets were collected and flash-frozen for future protein purification.

##### Protein constructs

BSA (Fisher: 23209) and Tubulin (PurSolutions LLC: 032005) are bovine versions and proteins were acquired directly from vendors. Tubulin was labeled with commercial NHS-conjugated dye (Cy5) according to supplier’s instructions (Sigma: GEPA150101) and its active labeled version purified using a published protocol (1). All remaining proteins are *Xenopus laevis* versions. DNA sequences were sourced from in-house plasmids, the *Xenopus laevis* Gene Collection (Source Biosciences), or synthesized (GenScript).

All TXP2 constructs were cloned as N-terminally tagged Strep6xHisGFP-TEV-TPX2 fusions using a modified pST50 vector (2) and cloned via Gibson assembly (NEB: E2611L). The same vector and cloning strategy were used to generate EB1-mCherry6xHis (3)(4) and His-GFP-PreScission-importin-β. GST-PreScission-importin-α and GST-PreScission-importin-αΔIBB were cloned into a pGEX6P1 vector using Gibson cloning. Insert fragments were PCR amplified from plasmids containing the indicated gene, and in most cases, unmodified from its wild type sequence. Exceptions are: TPX2-DM (StrHisGFP-TEV-DM_TPX2 with K284A, K285A, K327A, K330A), which was custom synthesized (Genscript), and ΔNLS-TPX2 (StrHisGFP-TEV-TM_TPX2 (aa1-716 with K123A, K125A, K126A, K284A, K285A, K327A, K330A) and TPX2 aa1-178Δ123 (StrHisGFP-TEV-NT_1-178_TPX2 (N-Terminal aa1-178, with K123A, K125A, K126A), which were both generated using site directed mutagenesis (Q5^®^ Site-Directed Mutagenesis Kit, NEB, E0554S). All constructs were fully sequenced and confirmed to have no errors.

##### Protein purification

For all TPX2 constructs, a previously published purification scheme was used (5). Briefly, cells were lysed using an EmulsiFlex (Avestin) in lysis buffer (0.05M Tris-HCl, 0.015M Imidazole, 0.75M NaCl, pH 7.75) containing 0.0002M phenylmethylsulfonyl fluoride (PMSF), 0.006M β-mercaptoethanol (βME), cOmplete™ EDTA-free Protease Inhibitor tablet (Sigma 5056489001), and 1000U DNase I (Sigma 04716728001). Lysate was clarified, bound to Ni-NTA agarose beads (Qiagen 1018236), washed and eluted in lysis buffer containing 200mM Imidazole. Protein was further purified via gel filtration (Superdex 200 HiLoad 16/600, GE Healthcare – 28-9893-35) in CSF-XB buffer (0.01 M Hepes, 0.002M MgCl_2_, 0.0001M CaCl_2_, 0.004M Ethylene glycol-bis(2-aminoethylether)-N,N,N’,N’-tetra-acetic acid (EGTA), 10% w/v sucrose, pH-7.75) containing either 0.1 M KCl for extract assays or 0.5M KCl for condensate assays.

GST-importin-α, clarified lysates were prepared in the same way with the exception of the lysis buffer (0.05M Na_2_HPO_4_/NaH_2_PO_4_, 0.5M NaCl, 5 mM β-mercaptoethanol, pH=7.45) and they were bound to a GST affinity column (GSTrap™ Fast Flow, GE Healthcare: 17-5131-02). The column was washed (0.05M Na_2_HPO_4_/NaH_2_PO_4_, 0.5M NaCl, 5 mM β-mercaptoethanol, pH=7.45), protein was eluted (0.05M Na_2_HPO_4_/NaH_2_PO_4_, 0.5M NaCl, 5 mM β-mercaptoethanol, 100 mM Glutathione-reduced, pH=7.45) and peak fractions were pooled.

His-GFP-PreScission-importin-β cells were lysed as explained above (Binding buffer: 50 mM Tris, 750 mM NaCl, 10 mM β-mercaptoethanol, 2% glycerol, pH = 7.88) and spun at 27000 rpm for 30 minutes. The protein was further purified from the lysed supernatant via an NiNTA affinity purification using a His-Prep FF 16/10 column (Column Volume ~ 20 ml). The column was then washed with 10 column volumes (CV) of binding buffer and the bound importin-β was eluted with sharp Imidazole gradient (4 CV). The eluted fractions (~25 ml) were concentrated using an Amicon 50 kDa membrane to 10 ml. 500 μl of 6xHis-HRV-3C protease at 2 mg/ml were added to 10 ml protein solution (1:50 w/w; importin-β concentration was measured by absorbance at 280 nm using Nanodrop and the protein purity after His-purification was estimated to be 90%) and the sample dialyzed with a 20 kD cut-off against 4L of 25 mM HEPES, 150 mM NaCl, 5 mM β-mercaptoethanol, 5% glycerol, pH = 7.57.

After at least 16 h, the protein was taken out, concentrated to 4-5ml, use for size-exclusion chromatography (SEC) on a HiLoad Superdex 200-increase column (16/600) equilibrated in buffer (50 mM Tris, 100 mM NaCl, 10 mM β-mercaptoethanol, pH = 7.5). Fractions containing cleaved importin-β were pooled and concentrated to 500 ul, which was subjected to a final affinity purification to remove the protease in a His-Trap high performance (5 ml) column. The flow through with purity >96% was collected and concentrated to 100 mg/ml in CSF-XB containing 0.1M KCl and 10% Sucrose (pH = 7.85).

EB1-6xHis-mCherry was purified as described previously (4). All proteins were aliquoted into single-use volumes, flash frozen and stored at −80°C. Before use, all proteins were thawed on ice and pre-cleared of aggregates via ultracentrifugation at 80,000 RPM for 15min in a TLA100 rotor in an Optima MAX-XP ultracentrifuge at 4°C. Protein concentrations were determined by OD280 absorbance measured via nanodrop (model ND-1000) using the extinction coefficient of the proteins (6).

##### Size-Exclusion chromatography in line with Multi-Angle Light Scattering (SEC-MALS)

To achieve highest resolution and separation of protein complexes, Superdex-200-increase-3.2/300 was equilibrated in low salt CSF buffer with no crowder (0.01M HEPES, 0.002M MgCl_2_, 0.0001M CaCl_2_, 0.004M Ethylene glycol-bis(2-aminoethylether)-N,N,N’,N’-tetraacetic acid (EGTA), 6 mM β-mercaptoethanol, 0.1M KCl, pH-7.75) in line with a Wyatt scattering instrument operating with a 632.38 nm red laser and 18 detectors located every 20° to monitor scattered light at multiple angles. The flow rate was 0.04 ml/min and a 25 μl sample loop was used for injection. The molecular weights of eluted complex were further characterized by a Debye plot using Kc/R(θ) = 1/M_w_ + B_2_c (7). Prior to injection into the column, all the buffers and protein solutions were filtered with 0.22μm PTFE filters (hold up volume <10 μl, product number SLGV004SL from Milipore Sigma). 60 μM stock TPX2 in high salt CSF-XB containing 0.5 M KCl and 10% Sucrose was diluted to final of 2 μM in low salt CSF buffer, containing 0.1 M KCl and no sucrose, and the sample (final salt of 0.17M KCl) was injected to the column. To monitor the association of TPX2 to importins, the GFP-TPX2 concentration was fixed at 2μM and importins were added to a final concentration of 20μM (final salt of 0.17M KCl) prior to sample injection. Control samples containing only importins were injected at 20 μM in CSF containing 0.1M KCl and no sucrose.

##### Analytical Size Exclusion to test reversibility of TPX2 importin-β Complex

To test the reversibility of TPX2-importin-β complex, we conducted size exclusion chromatography with a constant TPX2 concentration of 2 μM while varying importin-β concentration ranging from 2 to 30 μM. A Superdex200-increase-3.2/300 column was equilibrated in CSF buffer containing 0.1 KCl, 6 mM β-mercaptoethanol and no sucrose, and the sample was injected as described above.

##### Biolayer interferometry (OCTET)

Binding kinetic measurements were performed using biolayer interferometry via an Octet instrument. Anti-Penta-His (HIS1K) sensors were purchased from ForteBio. The sensors were washed in CSF containing 0.5M KCl and no sucrose. The TPX2 or TPX2 fragments containing 6xHis-tags were loaded into sensor for 30-45 seconds at low concentration of 120 nM to ensure no aggregation occurs on the sensor surface. The sensor was then washed in low salt CSF containing 0.1 M KCl and no sucrose for 300 seconds prior to exposing to importin solutions. Equilibrium binding was performed in wells with varying concentrations of importins, for 900 seconds to ensure the binding curve was plateaued. At longer time scales, the association and dissociation rate of ligand reaches the equilibrium. Using a simple titration curve, the amplitude can be plotted as function of ligand concentration. Upon equilibrium, the response follows the equation *R = R_max_C*/(*K_d_ + C*),where R_max_ is the maximum plateau, and K_d_ is the equilibrium dissociation constant (8). The height of binding amplitude was obtained by subtracting the signal from the control sample with no importin. It was essential to include a control sample for every run, as the loading of TPX2 constructs on the sensor can vary. Each data was replicated in two distinct measurements and the errors were calculated. For importin-β binding assays, 0.01% (v/v) tween-20 was used in the buffer to eliminate non-specific binding to sensors. All the importin-α-ΔIBB data were collected in regular CSF buffer containing 0.1 M KCl, no tween, and no sucrose.

##### Condensation (phase separation) assay

Proteins were diluted to reach a 5x final concentration in a CSF-XB Buffer containing 500 mM KCl salt at 4°C, then diluted 1:4 in CSF-XB containing no salt at room temperature (23°C) to reach a final salt concentration of 100mM. To image condensates via epifluorescence microscopy, the reaction mixture was immediately pipetted into a flow chamber (constructed with double-stick tape, a glass slide and a 22×22mm coverslip). The slide was placed coverslip-side down into a humidity chamber for 10 minutes at room temperature to allow condensates to settle; reaction was then imaged via epi-fluorescent microscopy. Crowding agents were never used.

##### Static and Dynamic Light Scattering (SLS and DLS)

Light scattering measurements were performed with a Wyatt instrument operating with red laser (λ = 632.8 nm), with the detector fixed at 90° to monitor the scattered intensity. Protein samples were filtered with 0.22 μm filters (hold up volume <10ul) prior to measurements. Normalized intensity-intensity correlation functions *g*_2_(*q*,τ) of 20 second duration were recorded. We fitted the intensity-intensity correlation functions possessing one broad decaying exponent using the cumulant function (9):

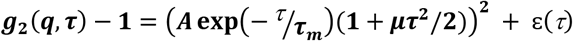

where *A* is the scattering amplitude, ε(***τ***) accounts for any noise in the signal, and ***μτ***^2^ is the polydispersity index. In our measurements, ε was one or two orders of magnitude lower than the amplitudes. For ***g***_2_s possessing two exponent decays, we fitted the data with two relaxation times corresponding to monomer/oligomers and condensates, respectively.

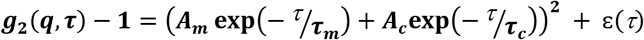

We used ***τ_m_*** to determine the average diffusivity, *D_m_*, from *D_m_* = (*q*^2^***τ***)^-1^, where 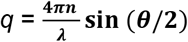 is the wave vector at a scattering angle of 90°, λ = 632.8 nm is the wavelength of the incident red laser and *n* = 1.33 is the solution refractive index.

In addition, we tested the same samples with DLS to monitor the importin ratio at which no sub-diffraction condensate was detected.

#### *Xenopus laevis* egg cytosol assays

##### Cytosol preparation

*Xenopus laevis* cytosol naturally arrested in meiosis II was prepared as described in (10). Briefly, *Xenopus laevis* eggs were collected after an overnight laying period. Eggs from individual frogs were kept separate but prepared in parallel, and typically 2 batches of eggs were used. Egg jelly coats were removed, and cytosol was fractionated away from egg yolk, membranes, nucleus and organelles by centrifugation (10200 RPM in HB-6 for 15 minutes). Eggs were constantly maintained at 18°C via preparation in a temperature-controlled room. Undiluted cytosol was collected, supplemented with Cytochalasin-D, protease inhibitors, ATP, and creatine phosphate and kept at 4°C until use.

*Xenopus* cytosol was immunodepleted of endogenous TPX2 as described in (5). Briefly, immunoaffinity purified antibodies against TPX2 or an unspecific IgG control antibody were conjugated to magnetic Dynabeads Protein A (ThermoFisher: 1002D) at 4°C overnight. Antibody-conjugated beads were split into two equal volume aliquots; supernatant was removed from one aliquot using a magnetic block and *Xenopus* cytosol was added. Beads were gently suspended in cytosol every 10 minutes for 40 minutes. Cytosol was removed from beads (using magnetic block) and then subjected to another round of depletion using the same procedure with the second aliquot of antibody-conjugated beads. Immunodepletion was assessed via Western blots and functional assays.

Branching MT nucleation assay: MT nucleation reactions were carried out as described in (5). Briefly, *Xenopus* cytosol was supplemented with fluorescently labeled tubulin ([1 μM] final) to visualize MTs, mCherry-fused End Binding protein 1 (EB1) ([100 nM] final) to track MT plus ends, and sodium orthovanadate ([0.5 μM] final) to inhibit dynein-mediated MT gliding. Mono-dispersed purified TPX2 (with or with our importins) was added at specified concentrations and excess molar ratios. In all experiments CSF-XB buffer containing 0.1 M KCl was used to match total dilution across all experiments (25% of extract volume). All reagents used were in CSF-XB buffer containing 0.1 M KCl. The reaction mixture was prepared on ice, then pipetted into a coverslip flow chamber at 18°C, which marked the start of the reaction.

Each reaction was imaged via TIRF microscopy for 30-40 minutes at 10-20 second intervals. Typically, two reactions (e.g. 0x and 1x excess importin-β) were prepared and flown into parallel chambers on to single slide, and each was imaged every 10-20 seconds by alternating between fields using the ‘XY acquisition’ function in NIS Elements (Nikon). This procedure enabled assessing multiple reactions within single cytosol preparation that has a finite lifetime (~2-6 hours). All experiments using *Xenopus* egg cytosol were reproduced at least three times using separate cytosol preparations. Similar results were seen in all replicates.

##### Analysis of branching MT nucleation

Total MT number per reaction was determined by counting the number of EB1 spots on MT plus ends in the entire field of view. EB1 detection was achieved via the ‘Color Threshold’ function (Otsu threshold) and the ‘Particle Analyzer’ function (size 0.1 −1 μm^2^) on FIJI. Parameters were optimized for each data-set according to visual assessment of tracking accuracy. Plots were generated by plotting the normalized number of EB1 detections (relative to 0x importins condition) for the 20-minute frame for each reaction.

Image collection and processing: The imaging technique used is indicated in each corresponding figure legend. Total Internal Reflection Fluorescence (TIRF) and Epifluorescence (Epi), microscopy methods were carried out on a Nikon TiE microscope with a 100X, 1.49NA oil immersion objective and an Andor Zyla sCMOS camera. All experiments were carried out at 18°C in a temperature-controlled room.

NIS-Elements software was used for all image acquisition. All images within a data set were taken with identical imaging parameters. Binning (2×2) was used in the case of *Xenopus* cytosol branching MT nucleation assays, but not for phase separation assays.

FIJI was used for all image analysis. Images of branching MT nucleation are 500 by 500 pixel crops of the center of the imaging field (comprising ¼ the area of the total field). Images of GFP-TPX2 phase separation are 100 by 100 pixel crops of a representative field. All images were processed using optimal brightness and contrast windows for each field to allow visualization of the relevant structure. Max/min windows between samples did not deviate more than 2-fold. Images within a figure panel were acquired and processed in the same setting.

Adobe illustrator/Inkscape was used to generate illustrations and compile figures. Oracle were used to generate all plots. The surface electrostatic potential and surface hydrophobic residues were visualized by Pymol for importin-β (PDB: 1QGK) and importin-αΔIBB (PDB: 4U5L).

**Fig. S1.**
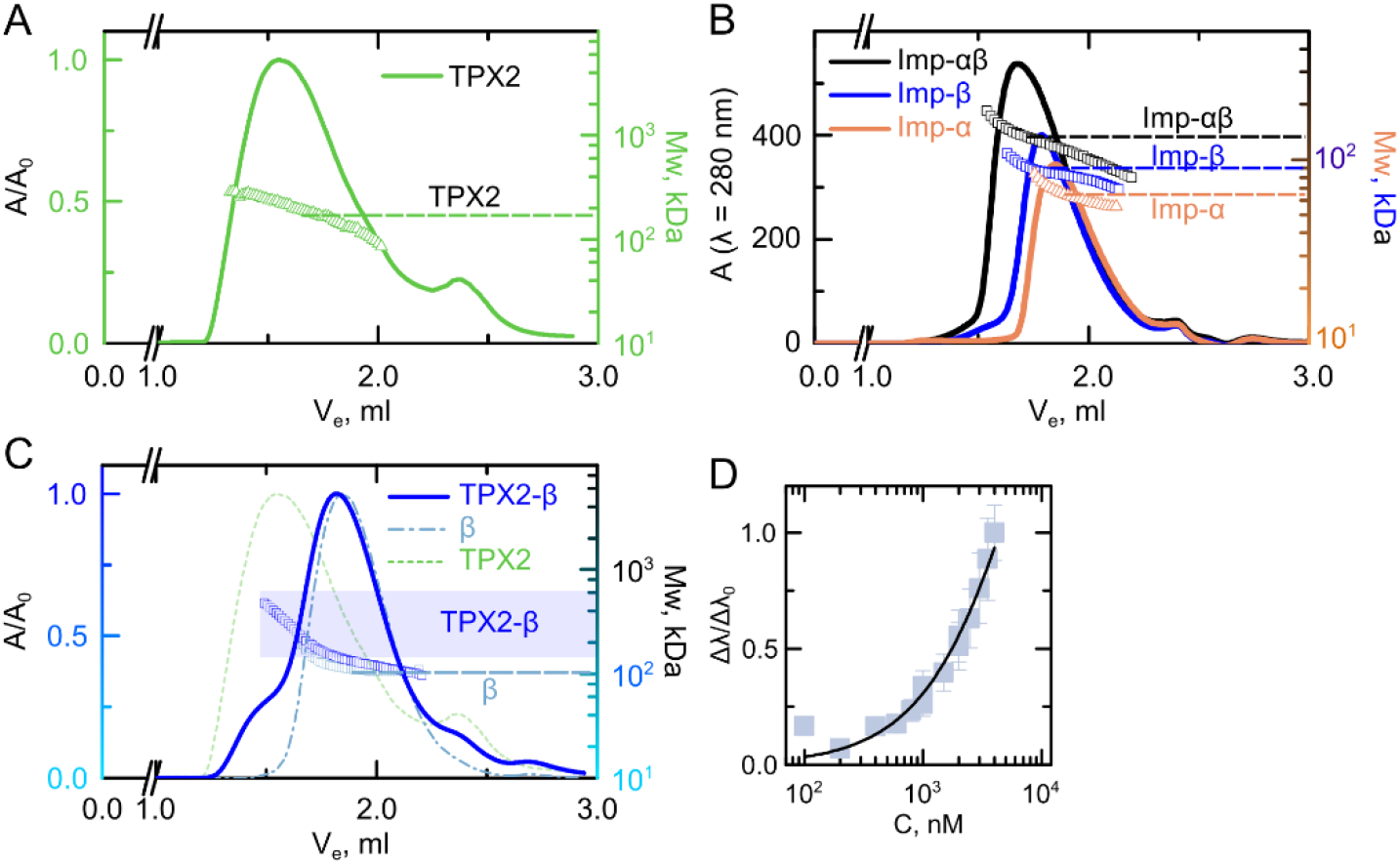
Characterization of importin-α/β heterodimer and TPX2-importin-β complex. (*A*) SEC-MALS elution profile of GFP-TPX2. The MALS profile indicates that GFP-TPX2 is monodisperse and mainly contains monomers. (*B*) SEC-MALS elution profile of GST-FL-importin-α (light brown), importin-β (blue), and GST-importin-α:importin-β (black). The arrows indicate the molecular weights of 63.6 kDa, 83.7 kDa, 130.0 kDa for GST-importin-α, importin-β, and GST-importin-α:importin-β respectively. (*C*) SEC-MALS elution profile of TPX2-importin-β complex (dark blue) and importin-β alone (light dashed blue). TPX2:importin-β MALS profile suggests there is a population of oligomers eluting from 1.2-1.6ml with molecular weights ranging from 200-700 kDa. (*D*) Normalized BLI-Octet binding curve for CT-TPX2 (319-716) and importin-β.

**Fig. S2.**
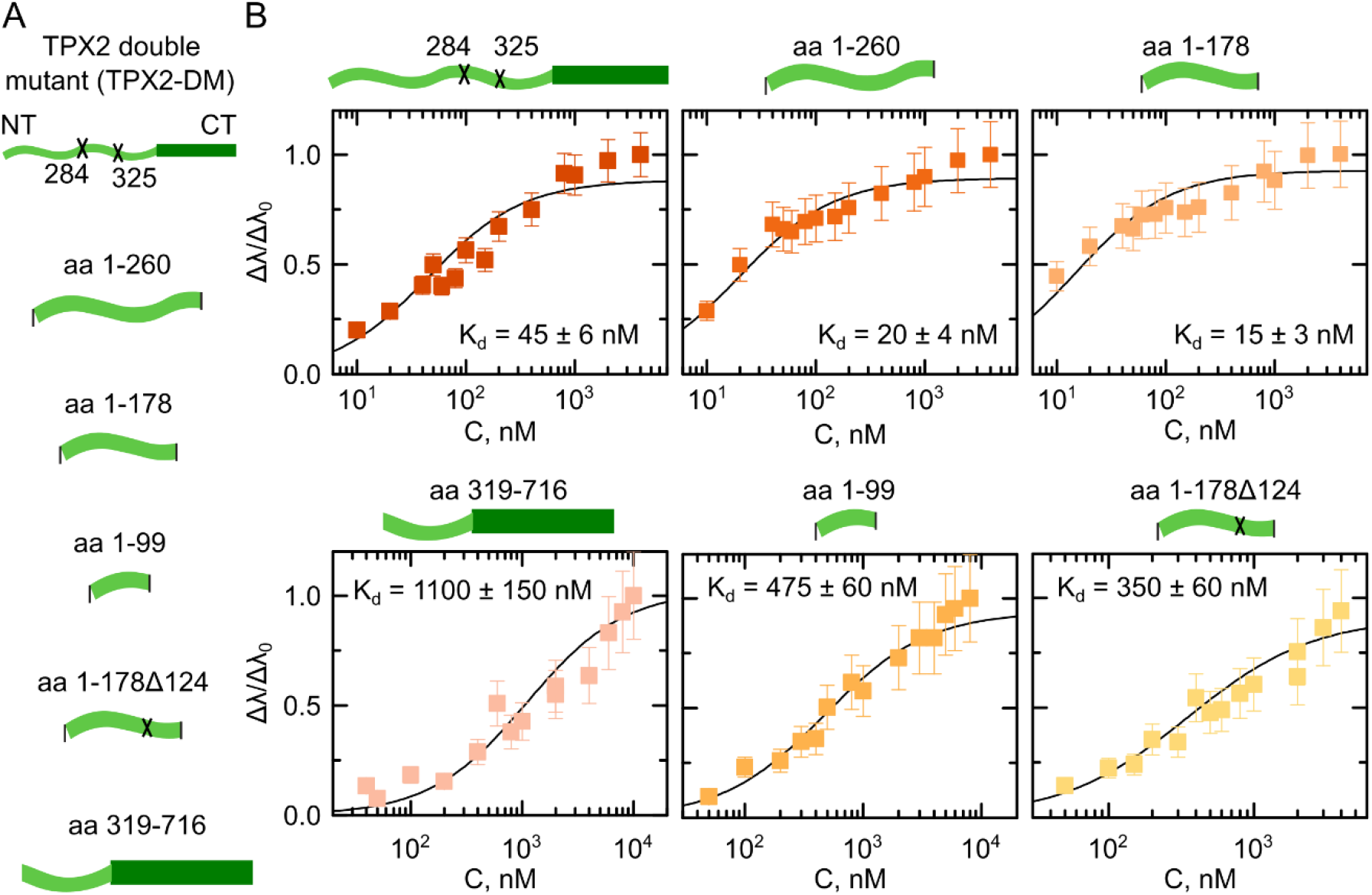
Identification of TPX2 putative NLS site aa123-126. (*A*) Schematics of tested TPX2 constructs for binding to importin-αΔIBB (*B*) Bio-layer interferometry binding curves for indicated TPX2 constructs with importin-αΔIBB.

**Fig. S3.**
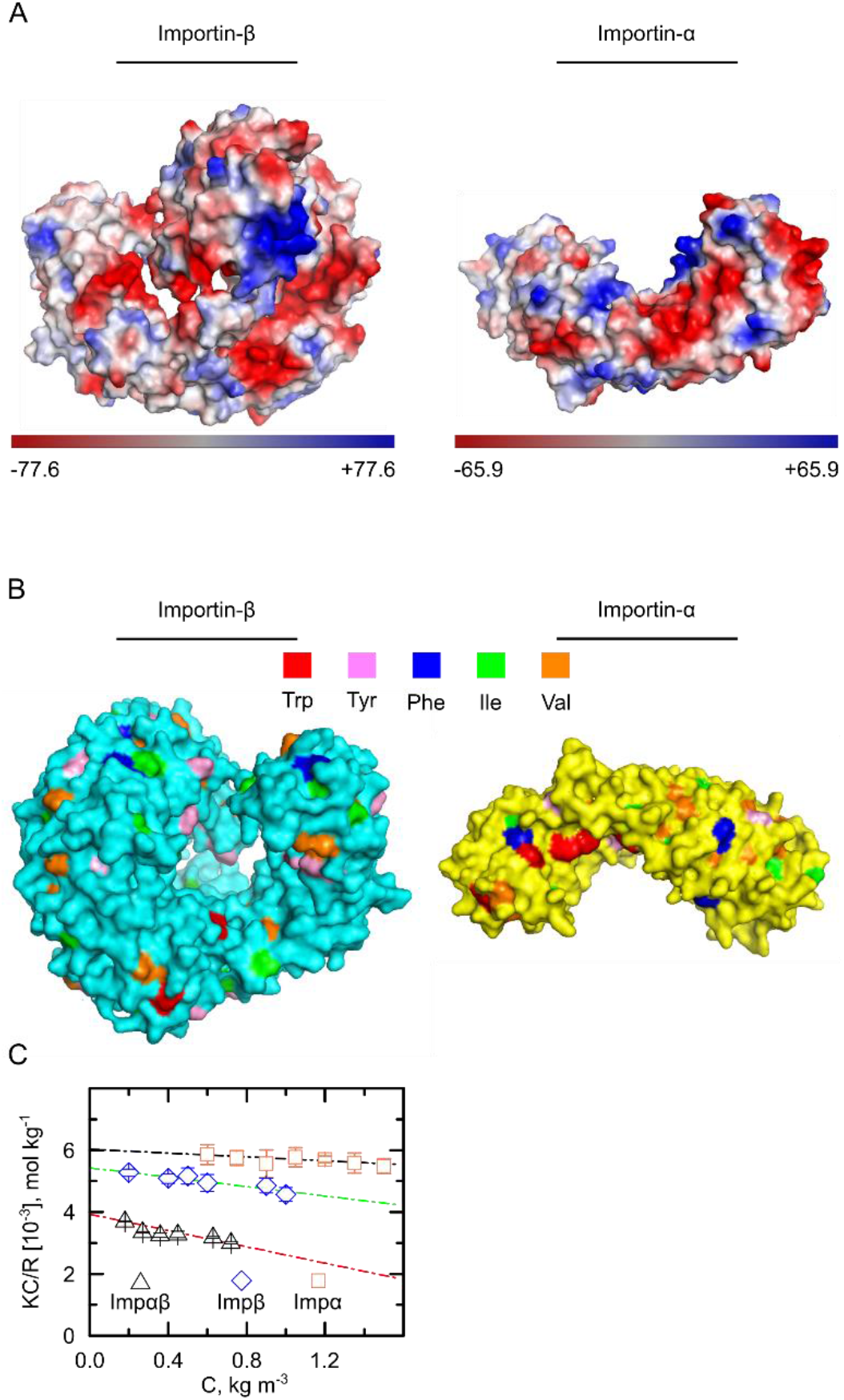
Illustration of exposed electrostatic potential and hydrophobic patches on importin surfaces. (*A*) Surface electrostatic potential of importin-β and importin-αΔIBB. The net charge of Importin-β and importin-αΔIBB are −38 and −9 respectively. The red and blue indicate negative and positive patches, respectively. (*B*) Exposed hydrophobic patches on the surface of importins. The residues are colored as Tryptophan (red), Tyrosine (pink), Phenylalanine (blue), Iso-leucine (green), and Valine (orange). (*C*) The osmotic compressibility of the importin.

